# nosoi: a stochastic agent-based transmission chain simulation framework in R

**DOI:** 10.1101/2020.03.03.973107

**Authors:** Sebastian Lequime, Paul Bastide, Simon Dellicour, Philippe Lemey, Guy Baele

## Abstract

The transmission process of an infectious agent creates a connected chain of hosts linked by transmission events, known as a transmission chain. Reconstructing transmission chains remains a challenging endeavor, except in rare cases characterized by intense surveillance and epidemiological inquiry. Inference frameworks attempt to estimate or approximate these transmission chains but the accuracy and validity of such methods generally lack formal assessment on datasets for which the actual transmission chain was observed. We here introduce `nosoi`, an open-source R package that offers a complete, tunable, and expandable agent-based framework to simulate transmission chains under a wide range of epidemiological scenarios for single-host and dual-host epidemics. `nosoi` is accessible through GitHub and CRAN, and is accompanied by extensive documentation, providing help and practical examples to assist users in setting up their own simulations. Once infected, each host or agent can undergo a series of events during each time step, such as moving (between locations) or transmitting the infection, all of these being driven by user-specified rules or data, such as travel patterns between locations. `nosoi` is able to generate a multitude of epidemic scenarios, that can – for example – be used to validate a wide range of reconstruction methods, including epidemic modeling and phylodynamic analyses. `nosoi` also offers a comprehensive framework to leverage empirically acquired data, allowing the user to explore how variations in parameters can affect epidemic potential. Aside from research questions, `nosoi` can provide lecturers with a complete teaching tool to offer students a handson exploration of the dynamics of epidemiological processes and the factors that impact it. Because the package does not rely on mathematical formalism but uses a more intuitive algorithmic approach, even extensive changes of the entire model can be easily and quickly implemented.

---

Infectious disease events, especially those resulting from novel emerging pathogens, have significantly increased over the past few decades, possibly as a result of alterations in various environmental, biological, socioeconomic, and political factors [1]. By definition, infectious agents need to spread through transmission between hosts. If successful, the resulting transmission process creates a connected chain of hosts linked by transmission events, usually called a transmission chain. Transmission is highly stochastic and can be influenced by a wide array of intrinsic and extrinsic factors, such as within-host dynamics and environmental or host behavioral factors. Reconstruction of transmission chains, however, remains difficult to achieve, except in certain rare cases characterized by intense surveillance and epidemiological inquiry [2, 3].

Molecular data may represent a critical asset in reconstructing the transmission history of a pathogen [3–7]. Often, however, the relationship between individual cases is too distant to allow for the perfect reconstruction of a transmission chain. In that context, the study of infectious agents’ genomic sequences can be used to reconstruct, under an evolutionary model, their likely evolutionary history. These reconstructions rely on evolution occurring on the same time scale as the epidemic or transmission process, which is the case for most fast-evolving pathogens such as RNA viruses [8, 9]. The inferred evolutionary history has been used in recent years to estimate the timing, the origin, or the effectiveness of mitigation measures of several epidemics [10–13].

The accuracy, validity, or limitations of both currently available and future methods, however, generally lack formal assessment on datasets for which we have been able to observe the actual geographical spread and the complex factors that shaped its pattern. In that context, a simulated dataset is extremely useful as the exact transmission history is known and can be compared to the histories inferred from different software packages. The last decade has seen the development of several integrated epidemic and genetic simulation tools that can be used to assess the performance of some of these models, such as TreeSim [14], SEEDY [15], outbreaker2 [16], or FAVITES [17].

While undoubtedly useful, these tools fall short in accommodating a wide range of epidemiological scenarios. In particular arboviral (e.g., Zika, dengue, or yellow fever) outbreaks, where two types of hosts participate in the epidemic process, are poorly modeled. These hosts are characterized by drastically different behavior or infection dynamics and cannot be accurately modeled using a single host type. Furthermore, geographic location diffusion is simulated in these tools, when possible, on a contact network or in discrete space. Yet recent years have seen the development of methods taking advantage of phylogeographic diffusion in continuous space [18, 19], creating a need for epidemiological simulations in a continuous space. To enable the performance assessment of these methods under complex and realistic scenarios, including spread in continuous space or arbovirus outbreaks, we present `nosoi`, a flexible agent-based transmission chain simulator implemented as an open-source R package [20].

## Characteristics

`nosoi` generalizes and significantly extends a basic model that allowed individual humans and mosquitoes – each one being characterized by a unique set of infection parameters – to interact within a simulated environment [21]. It was initially designed to model real-world arboviral epidemics unfolding under varying within-host dynamics [21].

`nosoi` employs agent-based modeling, which focuses on the individual active entities – known as (autonomous) agents – of a system and defines their behavior and the interactions between them. The main interest then lies in the global dynamics of and the complex phenomena within the system that emerges from the interactions of the many individual behaviors. Within `nosoi`, the agents’ behavior is governed by user-specified rules that can accommodate high levels of stochasticity at each level of the epidemic process. Agents can experience dual-host dynamics, such as those from human and mosquito populations, and exist in structured populations, with different behaviors according to host type and/or structure. Population structure can either be absent, discrete (e.g., different categories), or continuous (such as geographic space). In these structures, agents can trigger a movement, a contact, or a transmission event, with the probability of such an event occurring being potentially host-, individual-, structure- and/or time-dependent. These agents are recruited when infected and can either recover or die from the infection, resulting in their removal from the simulation. The status and location of each agent are assessed according to the model during each step of the discretized time of the simulation (Figure 1). The simulation ends when the user-specified value of the number of infected agents or when the targeted simulation time is reached.

**Figure 1.**
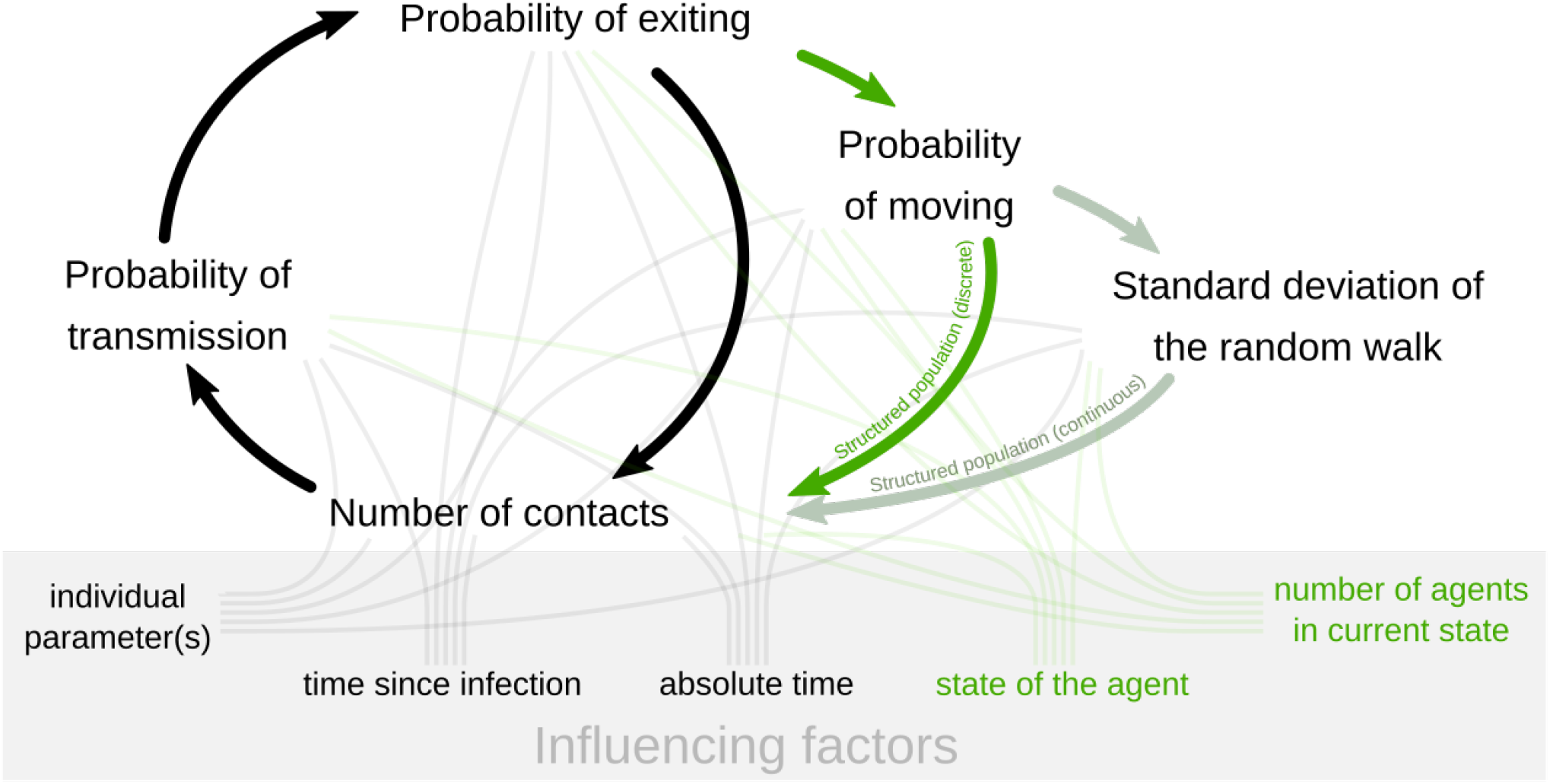
Schematic of status and location assessment for each agent (in case of a structured population), or host, during each discretized time step of the simulation. Optional steps in the simulation framework are shown in shades of green and are only performed in case of a structured (either discrete or continuous) population. Several factors (embedded in the gray box), either individually or globally set, can influence these steps according to user-specified settings.

In essence, `nosoi` allows the user to simulate and keep track of one or more transmission chains occurring during an infectious disease outbreak and, as such, to store and output a (collection of) transmission tree(s). Genetic data can be subsequently simulated along each transmission tree using sequence simulation software such as *π*buss [22] or SantaSim [23], which can then serve as input for phylodynamic inference methods. `nosoi` is accompanied by extensive tutorials, helping the user to set up and visualize their simulation, available as documentation in the package, or at https://slequime.github.io/nosoi/.

## Practical example

We here showcase `nosoi` with the starting scenario of a single human infected with an Ebolavirus-like pathogen in West Africa. The simulated epidemic unfolds in a geographically structured host population, specifically in a continuous geographic space, for 365 days or discrete time-steps. Within-host dynamics, influencing the probability of exiting the simulation (dying or recovering) and the between-host transmission probability, are modeled according to published literature that describes Ebolavirus infection in humans [24, 25]. The remaining parameters (number of daily contacts, probability of movement, and standard deviation of the random walk in continuous space) were empirically set. The number of daily contacts is restricted by the number of people living in the area, as provided by spatial demographics data obtained from WorldPop (www.worldpop.org), to avoid reaching locally unrealistic counts of infected humans. The complete specification and accompanying code for this simulation are available as a document on `nosoi`’s website (https://slequime.github.io/nosoi/articles/examples/ebola.html).

Over the course of 365 days, the simulation has yielded 3,603 infected agents. The average number of secondary cases per agent is 1.12, which is roughly coherent with previous epidemiological estimates of *R*_0_ for previous Ebolavirus outbreaks [26]. The increase in infected agents’ number is exponential, as would be expected considering the specifications of the model, i.e., absence of intervention strategies or changes in the simulated environment.

The transmission chain can be represented either as a network (Fig. 2A) or as a tree (Fig. 2B) that can be mapped in the continuous space in which the epidemic took place (Fig. 2C). The tree representation of the transmission chain can be seen as the genealogy of the pathogen population over which molecular evolution generates the observed sequence data, then used to reconstruct this same history. In this representation, each internal node is a transmission event, each tip represents the exit point in time of an agent, and the root is the starting point in time of the initially infected agent. Branches or sets of connected branches represent the life span of each agent. This tree is binary, counts as many tips as the total number of infected agents and as many internal nodes as transmission events.

**Figure 2.**
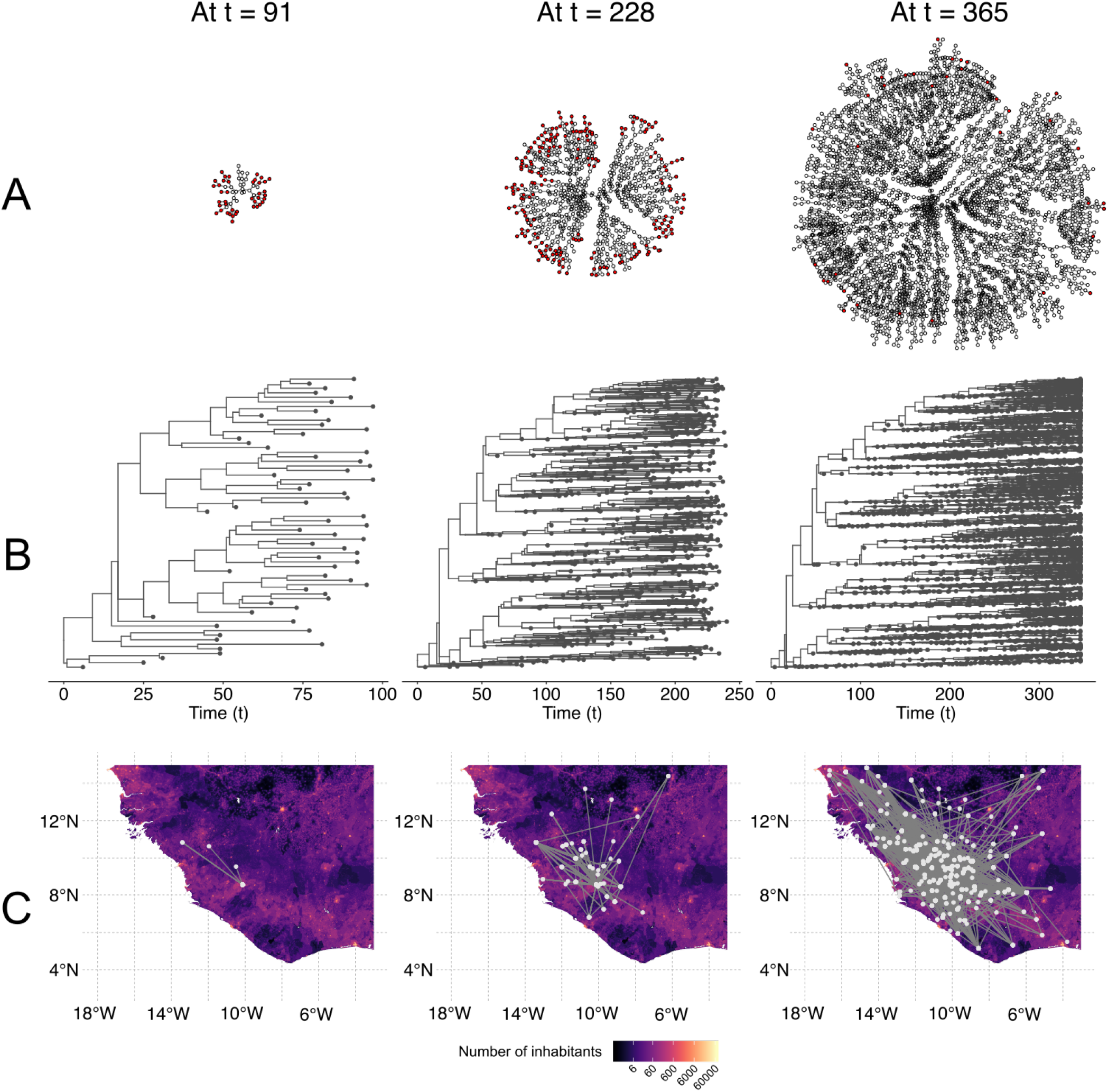
Visualization of a simulated Ebolavirus-like transmission chain in West Africa at three time-points (91, 228 and 365 days after the introduction of the first infected host), represented as (A) a network, (B) a tree or (C) a tree mapped on the continuous space the simulation took place in.

Other examples are available on `nosoi`’s website illustrating various scenarios, such as spread of a dengue-like pathogen (dual-host) in a discrete space or an unstructured population of hosts. The tutorials also provide guidelines on how to set up simulations in various combinations of settings currently available.

## Uses

Trends in globalization, including expansion in international travel and trade, have extended the reach and increased the pace at which infectious diseases spread [1]. These trends provide infectious agents with ample opportunities to establish and spread successfully, but many practical difficulties remain in accurately inferring key aspects of an epidemic. Standard testing of models of spread typically focuses on using that same model to generate simulated data, which offers important but limited insights and mostly provides a test of proper implementation and a way to compare different methodologies. `nosoi`, however, is a phylogenetic model-independent agent-based simulation framework that offers realistic and complex epidemiological scenarios. As such, it enables accurate testing of popular inference methods in both discrete and continuous phylogeography using either maximum-likelihood [27] or Bayesian inference [18,28,29], which are widely used in pathogen phylodynamics. In that regard, an interesting application of our proposed simulation framework could be to study the increasingly popular structured coalescent models [30–32], and to compare their accuracy under realistic epidemiological transmission scenarios against discrete phylogeographic inference.

`nosoi` enables the simulation of real-life scenarios of viral outbreaks, and we provide several example scenarios to showcase its capabilities to generate a single transmission chain using different settings. An important aspect is that the resulting transmission tree, which describes the transmission events between infected hosts, differs from the phylogenetic tree, which describes the ancestral genetic relationships between pathogens sampled from these hosts. In that regard, it is crucial to acknowledge the growing number of methods that infer either phylogenetic trees, transmission trees, or jointly estimate both (for an overview, we refer to Baele *et al.* [33]).

Apart from assessing the performance of various methods in reconstructing geographic spread or the dynamics of an infectious agent, `nosoi` can prove useful for assessing the performance of classic deterministic SIR and SIRS compartmental models [34]. These epidemiological models estimate the theoretical number of people infected with a contagious illness in a closed population over time under some assumptions. For example, the original SIR model assumes that the population size is fixed, that the incubation period of the infectious agent is instantaneous, and that the duration of infectivity is the same as the length of the disease. It also assumes a completely homogeneous population with no age, spatial, or social structure. These assumptions can be matched as closely as possible by the user-defined settings in `nosoi` or be violated in more realistic settings, allowing to examine the sensitivity of the deterministic models to the assumptions under a complex and fine-tuned epidemiological scenario.

`nosoi` also offers, in line with its initial purpose [21], a comprehensive framework to leverage empirically acquired data. A pathogen’s within-host dynamics characterized in lab settings can be embedded into a full stochastic epidemiological model, allowing the user to explore how variation can affect its epidemic potential.

Aside from research questions, `nosoi` can provide lecturers with a complete teaching tool to offer students a hands-on exploration of the dynamics of epidemiological processes and the factors that impact it. Because the package does not rely on mathematical formalism but uses a more intuitive algorithmic approach, even extensive changes of the entire model or part of it can be easily and quickly implemented. The documentation provides suggestions for visualization using well-known external R-packages, such as `ggplot2` [35] or `ggtree` [36, 37]. The package is also fully integrated in the R and phylogenetic environments, and, through the use of the `treeio` and `tidytree` R packages [38], simulated transmission trees can be exported in a wide variety of formats for downstream analyses, such as the BEAST [29] or jplace [39] formats.

In summary, `nosoi` provides a complete, tunable, and expandable framework to simulate epidemiological processes based on transmission chains, in a user-friendly manner. Accessible through GitHub and the CRAN, the code is well covered by unitary tests and accompanied by extensive documentation, providing help and practical examples to users. Open-source and coded in the widely used R language, it allows users to customize their model by implementing new mechanisms for all or part of the core model. In addition, and contrary to other available tools, by decoupling sequence evolution from the epidemiological process, it can connect to any external sequence simulator, allowing the user to choose a tool and model that can address the biological question of interest.

## Acknowledgments

The authors would like to thank Maude Jacquot and Albin Fontaine for conducting preliminary tests with the simulator, and Mandev Gill for insightful discussions. SL and PB are post-doctoral research fellows funded by the Fonds Wetenschappelijk Onderzoek (FWO, Belgium). SD is supported by the Fonds National de la Recherche Scientifique (FNRS, Belgium) and was previously funded by the Fonds Wetenschappelijk Onderzoek (FWO, Belgium). PL acknowledges support by the Research Foundation – Flanders (‘Fonds voor Wetenschappelijk Onderzoek – Vlaanderen’, G066215N, G0D5117N and G0B9317N). GB acknowledges support from the Interne Fondsen KU Leuven / Internal Funds KU Leuven under grant agreement C14/18/094, and the Research Foundation – Flanders (‘Fonds voor Wetenschappelijk Onderzoek – Vlaanderen’, G0E1420N). The research leading to these results has received funding from the European Research Council under the European Union’s Horizon 2020 research and innovation programme (grant agreement no. 725422-ReservoirDOCS). The Artic Network receives funding from the Wellcome Trust through project 206298/Z/17/Z.

## References

1. Chan, E. H. et al. Global capacity for emerging infectious disease detection. Proceedings of the National Academy of Sciences of the United States of America 107, 21701–21706 (2010). URL http://www.pnas.org/cgi/doi/10.1073/pnas.1006219107.

2. Mollentze, N. et al. A Bayesian approach for inferring the dynamics of partially observed endemic infectious diseases from space-time-genetic data. Proceedings of the Royal Society B-Biological Sciences 281, 20133251–20133251 (2014). URL http://rspb.royalsocietypublishing.org/cgi/doi/10.1098/rspb.2013.3251.

3. Worby, C. J. et al. Reconstructing transmission trees for communicable diseases using densely sampled genetic data. Annals of Applied Statistics 10, 395–417 (2016). URL http://projecteuclid.org/euclid.aoas/1458909921.

4. Didelot, X., Gardy, J. & Colijn, C. Bayesian Inference of Infectious Disease Transmission from Whole-Genome Sequence Data. Molecular biology and evolution 31, 1869–1879 (2014). URL https://academic.oup.com/mbe/article-lookup/doi/10.1093/molbev/msu121.

5. Didelot, X., Fraser, C., Gardy, J. & Colijn, C. Genomic Infectious Disease Epidemiology in Partially Sampled and Ongoing Outbreaks. Molecular biology and evolution 34, 997–1007 (2017). URL http://eutils.ncbi.nlm.nih.gov/entrez/eutils/elink.fcgi?dbfrom=pubmed&id=28100788&retmode=ref&cmd=prlinks.

6. De Maio, N., Worby, C. J., Wilson, D. J. & Stoesser, N. Bayesian reconstruction of transmission within outbreaks using genomic variants. PLoS computational biology 14, e1006117–23 (2018). URL http://dx.plos.org/10.1371/journal.pcbi.1006117.

7. Campbell, F., Cori, A., Ferguson, N. & Jombart, T. Bayesian inference of transmission chains using timing of symptoms, pathogen genomes and contact data. PLoS computational biology 15, e1006930 (2019). URL http://dx.plos.org/10.1371/journal.pcbi.1006930.

8. Ypma, R. J. F., van Ballegooijen, W. M. & Wallinga, J. Relating Phylogenetic Trees to Transmission Trees of Infectious Disease Outbreaks. Genetics 195, 1055–1062 (2013). URL http://www.genetics.org/lookup/doi/10.1534/genetics.113.154856.

9. Romero-Severson, E., Skar, H., Bulla, I., Albert, J. & Leitner, T. Timing and Order of Transmission Events Is Not Directly Reflected in a Pathogen Phylogeny. Molecular biology and evolution 31, 2472–2482 (2014). URL https://academic.oup.com/mbe/article-lookup/doi/10.1093/molbev/msu179.

10. Dellicour, S. et al. Phylodynamic assessment of intervention strategies for the West African Ebola virus outbreak. Nature Communications 9, 2222 (2018). URL http://www.nature.com/articles/s41467-018-03763-2.

11. Dudas, G., Carvalho, L. M., Rambaut, A. & Bedford, T. MERS-CoV spillover at the camel-human interface. eLife 7, 250 (2018). URL https://elifesciences.org/articles/31257.

12. Hill, S. C. et al. Emergence of the Asian lineage of Zika virus in Angola: an outbreak investigation. The Lancet. Infectious diseases 19, 1138–1147 (2019). URL https://linkinghub.elsevier.com/retrieve/pii/S1473309919302932.

13. Grubaugh, N. D. et al. Travel Surveillance and Genomics Uncover a Hidden Zika Outbreak during the Waning Epidemic. Cell 178, 1057–1071.e11 (2019). URL https://linkinghub.elsevier.com/retrieve/pii/S0092867419307834.

14. Stadler, T. & Bonhoeffer, S. Uncovering epidemiological dynamics in heterogeneous host populations using phylogenetic methods. Philosophical transactions of the Royal Society of London. Series B, Biological sciences 368, 20120198 (2013). URL https://royalsocietypublishing.org/doi/10.1098/rstb.2012.0198.

15. Worby, C. J. & Read, T. D. ‘SEEDY’ (Simulation of Evolutionary and Epidemiological Dynamics): An R Package to Follow Accumulation of Within-Host Mutation in Pathogens. Plos One 10, e0129745 (2015). URL http://dx.plos.org/10.1371/journal.pone.0129745.

16. Campbell, F. et al. outbreaker2: a modular platform for outbreak reconstruction. Bmc Bioinformatics 19, 320–8 (2018). URL https://bmcbioinformatics.biomedcentral.com/articles/10.1186/s12859-018-2330-z.

17. Moshiri, N., Ragonnet-Cronin, M., Wertheim, J. O. & Mirarab, S. FAVITES: simultaneous simulation of transmission networks, phylogenetic trees and sequences. Bioinformatics 35, 1852–1861 (2019). URL http://eutils.ncbi.nlm.nih.gov/entrez/eutils/elink.fcgi?dbfrom=pubmed&id=30395173&retmode=ref&cmd=prlinks.

18. Lemey, P., Rambaut, A., Welch, J. J. & Suchard, M. A. Phylogeography Takes a Relaxed Random Walk in Continuous Space and Time. Molecular biology and evolution 27, 1877–1885 (2010). URL http://mbe.oxfordjournals.org/cgi/doi/10.1093/molbev/msq067.

19. Dellicour, S., Rose, R., Faria, N. R., Lemey, P. & Pybus, O. G. SERAPHIM: studying environmental rasters and phylogenetically informed movements. Bioinformatics 32, 3204–3206 (2016). URL https://academic.oup.com/bioinformatics/article-lookup/doi/10.1093/bioinformatics/btw384.

20. R Core Team. R: A Language and Environment for Statistical Computing. Vienna, Austria (2019). URL https://www.R-project.org/.

21. Fontaine, A. et al. Epidemiological significance of dengue virus genetic variation in mosquito infection dynamics. PLoS Pathogens 14, e1007187–21 (2018). URL https://dx.plos.org/10.1371/journal.ppat.1007187.

22. Bielejec, F. et al. πBUSS: a parallel BEAST/BEAGLE utility for sequence simulation under complex evolutionary scenarios. Bmc Bioinformatics 15, 133 (2014). URL http://bmcbioinformatics.biomedcentral.com/articles/10.1186/1471-2105-15-133.

23. Jariani, A. et al. SANTA-SIM: simulating viral sequence evolution dynamics under selection and recombination. Virus Evolution 5, 301–8 (2019). URL https://academic.oup.com/ve/article/doi/10.1093/ve/vez003/5372481.

24. Casillas, A. M., Nyamathi, A. M., Sosa, A., Wilder, C. L. & Sands, H. A current review of Ebola virus: pathogenesis, clinical presentation, and diagnostic assessment. Biological research for nursing 4, 268–275 (2003). URL http://journals.sagepub.com/doi/10.1177/1099800403252603.

25. Skrip, L. A. et al. Characterizing risk of Ebola transmission based on frequency and type of case-contact exposures. Philosophical transactions of the Royal Society of London. Series B, Biological sciences 372, 20160301 (2017). URL https://royalsocietypublishing.org/doi/10.1098/rstb.2016.0301.

26. Van Kerkhove, M. D., Bento, A. I., Mills, H. L., Ferguson, N. M. & Donnelly, C. A. A review of epidemiological parameters from Ebola outbreaks to inform early public health decision-making. Scientific Data 2, 150019–10 (2015). URL http://www.nature.com/articles/sdata201519.

27. Ishikawa, S. A., Zhukova, A., Iwasaki, W. & Gascuel, O. A Fast Likelihood Method to Reconstruct and Visualize Ancestral Scenarios. Molecular biology and evolution 36, 2069–2085 (2019). URL https://academic.oup.com/mbe/article/36/9/2069/5498561.

28. Lemey, P., Rambaut, A., Drummond, A. J. & Suchard, M. A. Bayesian phylogeography finds its roots. PLoS computational biology 5, e1000520 (2009). URL http://dx.plos.org/10.1371/journal.pcbi.1000520.

29. Suchard, M. A. et al. Bayesian phylogenetic and phylodynamic data integration using BEAST 1.10. Virus Evolution 4, vey016 (2018). URL https://academic.oup.com/ve/article/doi/10.1093/ve/vey016/5035211.

30. De Maio, N., Wu, C.-H., O’Reilly, K. M. & Wilson, D. New Routes to Phylogeography: A Bayesian Structured Coalescent Approximation. PLoS Genet 11, e1005421 (2015). URL http://dx.plos.org/10.1371/journal.pgen.1005421.

31. Müller, N. F., Rasmussen, D. A. & Stadler, T. The Structured Coalescent and Its Approximations. Molecular biology and evolution 34, 2970–2981 (2017). URL https://academic.oup.com/mbe/article/34/11/2970/3896419.

32. Bouckaert, R. et al. BEAST 2.5: An advanced software platform for Bayesian evolutionary analysis. PLoS computational biology 15, e1006650 (2019). URL http://dx.plos.org/10.1371/journal.pcbi.1006650.

33. Baele, G., Suchard, M. A., Rambaut, A. & Lemey, P. Emerging Concepts of Data Integration in Pathogen Phylodynamics. Systematic biology 66, e47–e65 (2017). URL https://academic.oup.com/sysbio/article/doi/10.1093/sysbio/syw054/2670001/Emerging-concepts-of-data-integration-in-pathogen.

34. Kermack, W. O. & McKendrick, A. G. A Contribution to the Mathematical Theory of Epidemics. Proceedings of the Royal Society A: Mathematical, Physical and Engineering Sciences 115, 700–721 (1927). URL http://rspa.royalsocietypublishing.org/cgi/doi/10.1098/rspa.1927.0118.

35. Wickham, H. ggplot2. Elegant Graphics for Data Analysis (Springer New York, New York, NY, 2009). URL http://link.springer.com/10.1007/978-0-387-98141-3.

36. Yu, G., Smith, D. K., Zhu, H., Guan, Y. & Lam, T. T.-Y. ggtree: an rpackage for visualization and annotation of phylogenetic trees with their covariates and other associated data. Methods in Ecology and Evolution 8, 28–36 (2016). URL http://doi.wiley.com/10.1111/2041-210X.12628.

37. Yu, G., Lam, T. T.-Y., Zhu, H. & Guan, Y. Two Methods for Mapping and Visualizing Associated Data on Phylogeny Using Ggtree. Molecular biology and evolution 35, 3041–3043 (2018). URL https://academic.oup.com/mbe/article/35/12/3041/5142656.

38. Wang, L.-G. et al. treeio: an R package for phylogenetic tree input and output with richly annotated and associated data. Molecular biology and evolution 60, 291 (2019). URL https://academic.oup.com/mbe/advance-article/doi/10.1093/molbev/msz240/5601621.

39. Matsen, F. A., Hoffman, N. G., Gallagher, A. & Stamatakis, A. A format for phylogenetic placements. Plos One 7, e31009 (2012). URL https://dx.plos.org/10.1371/journal.pone.0031009.

